# Differential universal ortholog composition of *Coffea arabica* L. sub-genomes and its contribution to regulatory networks governing essential biological processes

**DOI:** 10.1101/2023.03.08.531780

**Authors:** Thales Henrique Cherubino Ribeiro, Raphael Ricon de Oliveira, Chalfun-Junior Antonio

## Abstract

The polyploidy of *Coffea arabica* is an important trait affecting the evolution of this species. Genetic variability is scarce due to its recent origin as an interspecific hybrid from a single successful crossing event between *Coffea canephora* and *Coffea eugenioides* relatives. To further investigate the genomic composition of an allotetraploid we coupled high-throughput methodologies of co-expression analysis and full-length protein coding genes inference. Many of the expected orthologs were found to be missing from one of the two homoeologous chromosomes. The gene expression machinery is mainly represented by single-copy essential orthologs located in the *Coffea eugenioides* sub-genome. This result suggests a preference of the transcriptional and RNA processing machinery to be regulated by one parental sub-genome. To understand the operational modules of the sub-genomes transcription, we performed co-expression analysis that revealed 23 co-regulated modules. This system-wide approach clarified how biological processes (i.e., photosynthesis, cell wall biogenesis, translation, transcription, catabolism and biosynthesis) are running in synchrony and reinforces that there is an ongoing selective pressure in *C. arabica* that constrains the number of copies of some universal orthologues. Thus, this work contributes to our understanding of genome evolution in recent polyploids and supports crop breeding programs.

## Introduction

*Coffea arabica* L. (Rubiaceae) is considered the crop with the lowest level of genetic diversity reported so far (Scalabrin et al. 2020). It is an autogamous species that arose after a single event of interspecific hybridization, becoming the single polyploid species of the *Coffea* genus (Charrier and Berthaud 1985; Davis et al. 2006). The few single nucleotide polymorphisms identified within *C. arabica* are not shared with any of its potential parental species, *Coffea canephora* L. and *Coffea eugenioides* Moore (Scalabrin et al. 2020). This finding suggested a severe genetic bottleneck effect once the variation found in *C. arabica* arose after its recent polyploidization event. In addition, no major introgressions could be verified (Scalabrin et al. 2020). To understand how genetic diversity impacts the coffee economy several initiatives to investigate this allopolyploidy genome-and relative species genomes-are undergoing to better understand its organization and underlying regulatory networks (Lashermes et al. 2016; Tran et al. 2018a, 2018b; Mekbib et al. 2022; Cenci et al. 2012; Denoeud et al. 2014).

A central concept in studying allopolyploids is homoeology that is defined as “genes or chromosomes in the same species that originated by speciation and were brought back together in the same genome by allopolyploidization” (Glover et al. 2016). Cytogenetic studies in *C. arabica* showed that large genomic duplications or deletions did not occur, confirming the low structural divergence of homoeologous chromosomes between the sub-genomes inherited from the two diploid progenitor species (Pinto-Maglio 2006). The only major interchange between the sub genomes seems to be a single instance of homoeologous replacement at the tip of chromosome 7 (Scalabrin et al. 2020). One of the tips of the inherited homoeologue from *C. canephora* was replaced with a 1.7 million bases stretch from its *C. eugenioides* counterpart in an apparent homology-directed repair of double strand break (Scalabrin et al. 2020).

To become a functional genome, with ability to both allow a coordinated metabolism and reproductive success, *C. arabica* chromosomes needed to survive the original hybridization event and overall genome organization until a stable version was achieved (Lashermes et al. 2016). Although the chromosomal structure is similar between the sub-genomes, bioinformatic analyses using k-mers of 51 bp suggested high levels of sequence diversity between both homoeologs (Scalabrin et al. 2020) and differential expression of homoeologous genes was verified (Vidal et al. 2010). In the long run, these sub-genome specific sequence divergences may be the reason why polyploid genomes tend to turn into diploids (Wolfe 2001).

The evolution of genomes after their duplication is, to some degree, governed by gene interaction networks (Koonin and Wolf 2006). Connected genes, such as those that form functional modules like ribosomes and transcriptional complexes, have a tendency of not being lost following tetraploidy because a co-expression imbalance may lead to reduced fitness (Papp et al. 2003). This Gene Balance Hypothesis (Freeling and Thomas 2006) also predicts that homoeologs that are not tightly co-regulate with other genes-they are not dosage sensitive-will eventually evolve to singletons due to purifying selection (Papp et al. 2003). These additional copies may become pseudogenes or evolve novel functions.

The functional modules underlying gene co-expression networks may be found in contiguous regions of reduction-resistant pairs (Sankoff et al. 2010). This can happen because the selective pressure to maintain the balance of dosage sensitivity loci constrains the loss of these duplicated genes. So, there is a tendency to cluster co-expressed modules on chromosomes (Freeling and Thomas 2006). In this work we took advantage of a publicly available chromosome level assembly of *C. arabica* (Johns Hopkins University 2018) to (I) predict and annotate protein coding genes (PCG), (II) distinguish the PCG content between sub-genomes and (III) identify co-expression modules to provide insights on leaf metabolism.

The *C. arabica* genome is probably in a phase between a recent whole genome duplication event and the reestablishment of a diploid state. We verified contrasting results of homoeolog gene loss and retention that may be explained within the realm of evolutionary systems biology by linking the evolution of genes to their function within networks (Koonin and Wolf 2006). Although both sub-genomes are working in synchrony, it is possible that the kernel of this operating system-the system that controls the execution and memory allocation of other codes-is running from the *C. eugenioides* sub-genome.

## Results

The accurate prediction of protein coding *loci* in genomes requires species-specific parameters for its underlying Hidden Markov Model (HMM) (Hoff and Stanke 2019). To define those parameters, we gathered *ab initio* and extrinsic evidence in order to improve the accuracy and completeness of the annotation. The extrinsic evidence was based on a set of 174 billion nucleotides sequenced in 81 RNAseq libraries of multiple coffee tissues summarized in Supplemental Table S1. Both single and paired end reads were mapped to a fasta file containing the sub-genomes and excluding the unmapped contigs-regions of the genome that are repetitive and difficult to define chromosome coordinates with accuracy. In addition, these RNAseq fragments were also used for the *de novo* transcriptome assembly and transcriptome-based inference to provide full-length transcript data as an additional extrinsic evidence source for model training. This data provided hints for the coordinates of exons, introns and UTRs in the assembled *C. arabica* chromosomes.

After training a supervised machine learning model AUGUSTUS predicted a total of 69,464 full-length putative PCG being 30,162 in the *C. canephora* sub-genome and 39,302 in the *C. eugenioides* sub-genome (Supplemental Figure S1). Approximately 4.6 thousand full-length genes are putative species-specific (orphans) and 16 thousand had their expression verified in leaves (Figure 1). The putative PCG sequences, coordinates and annotation are available at https://dbi.ufla.br/lfmp/ca_annotation.

**Figure 1.**
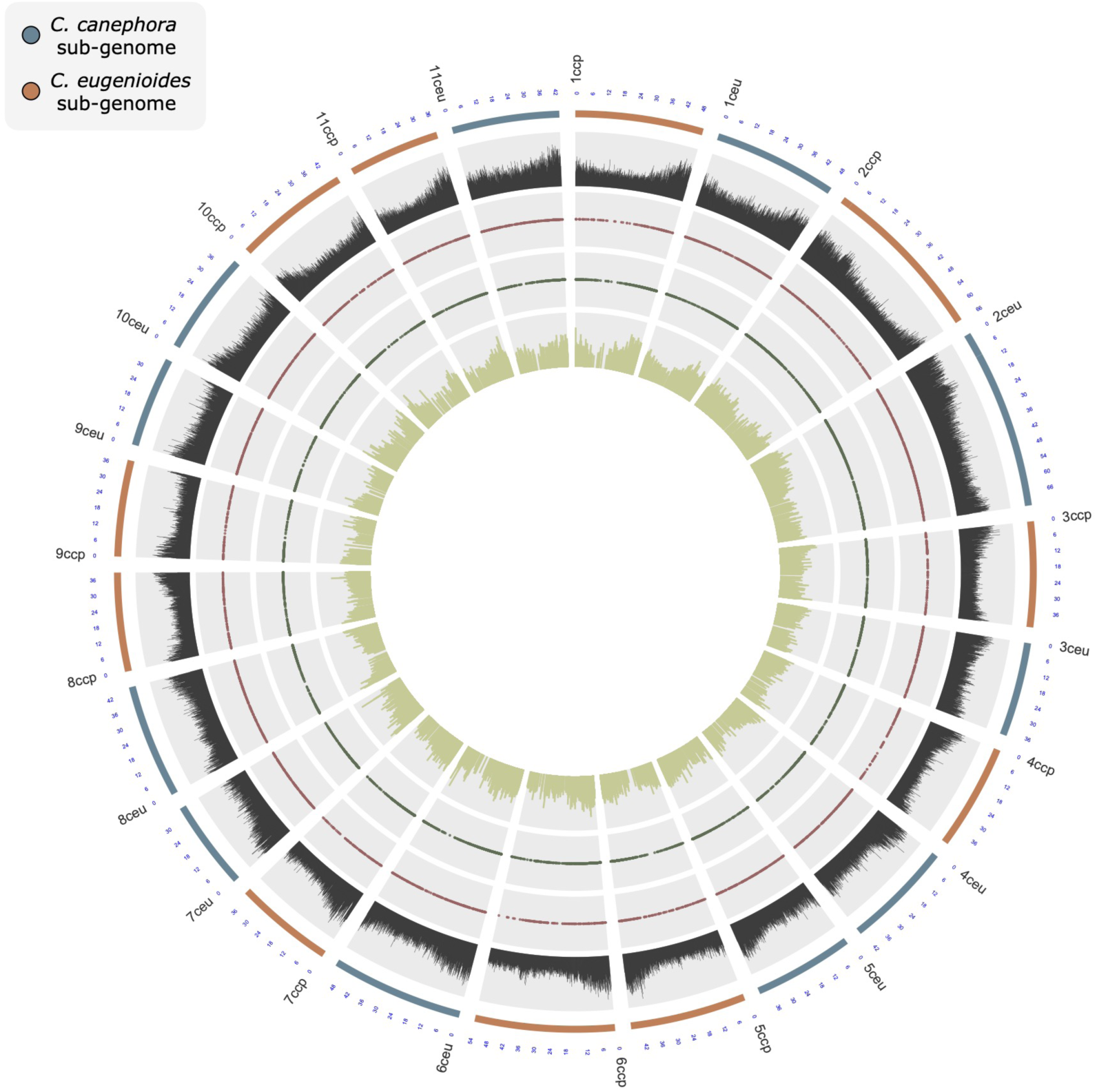
Genome-wide representation of Coffea arabica assembled homoeolog chromosomes. Concentrical cycles, left side; the outermost cycle represents chromosomes from each parental ancestor; from C. eugenioides (ceu) or C. canephora (ccp). First inner cycle; Inferred density of putative protein coding genes within a window of 0.1 million bases. Second inner cycle; red dots point out the chromosomal coordinates of potential Coffea arabica orphan genes (N = ∼4.6K). Third inner cycle; dark green dots point out the chromosomal coordinates of PCG expressed in leaves of the Experimental Group G (Table 1) (N=∼16K). Innermost cycle; bar plots depicting the sum of expression of all RNAseq samples from leaves. Expression values are reported in the logarithmic base two of counts per million (CPM).

### *De novo* transcriptome assembly and protein coding transcripts inference was an effective tool to retrieve BUSCO signatures

Because the short read alignment-based evidence does not distinguish non-coding from coding transcripts-the *RNAseq* methodology reads transcripts with poly-A tail-we decided to *de novo* assemble the transcriptome of each of the 6 groups of different sets of organs/experiments (Supplemental Table S1). Then, we filtered for the sequences with high probability of being transcribed into full-length proteins with the Transdecoder software.

The combination of the transcriptome assembly from all RNAseq libraries yielded a total of 2,518,967 transcripts. After redundancy filtering of sequences with more than 80% of similarity at the amino acid level we produced a set of 518,787 non-redundant transcripts with the average length of 491 bp, median length of 182 bp and a N50 value of 1,694. Then, the transdecoder software identified 105,598 potential complete proteins in the combined *C. arabica* transcriptome. BUSCO analyses showed that our transcriptome assembly presented 93.5% of the expected orthologs based on the expected composition of eudicots in the database eudicots_odb10 (version from 09/10/2020), which is in accordance with previous reports of gene predictions from a scaffold-level genome assembly that identified 92.4% BUSCO signatures in the “Bourbon Vermelho” variety (Scalabrin et al. 2020). Once we found that both the number of the full-length *de novo* predicted proteins and their BUSCO assessment were adequate, we used both the amino acid and mRNA sequences as hint sources for the model training and testing.

### Part of the identified BUSCO signatures and the number of PCG are different between the sub-genomes

To better understand the composition of the predicted PCG in the *C. arabica* sub-genomes we performed Gene Ontology (GO), InterPro domain identification and BUSCO analysis. Because our PCG prediction was performed to maximize the quantification of RNAseq fragments mapped to exons we did not allow multiple gene isoforms that potentially shared exons. In addition, our focus was to provide a reliable reference of protein coding genes in a General Feature Format (gff) file to allow *RNAseq* expression analyses of full length PCG. With that strategy we avoided quantifying other genic loci that are not translated.

We only allowed a single isoform of each PCG. In addition, only loci containing identifiable transcription start site (tss), start codon, exon(s), stop codon and transcription termination site (tts) were reported in the gff file. This approach allowed us to increase the number of quantified *RNAseq* reads that were uniquely mapped to exons by a factor of 63% when compared to running the htseq-count script with default parameters using the NCBI *Coffea arabica* Annotation Release 100 (2018).

Our procedure of inferring PCG separately for each sub-genome, instead of performing training and predicting steps in a genome-wide approach, allowed us to better understand the differences between them. Approximately 43% of the PCG were found in the *C. canephora* sub-genome with protein N50 of 987. Meanwhile, 57% of the PCG were identified in the *C. eugenioides* sub-genome with protein N50 of 777. We found these differences in the number of genes and their length intriguing, however the raw quantification of predicted PCG and their sizes cannot provide extensive insights into the actual composition and/or quality of genome wide inferences. Because of that, additional tools were applied to measure quantitatively the completeness using evolutionarily informed expectations of gene content (Simão et al. 2015).

Essential genes are significantly enriched in lineage-specific universal orthologs databases of model organisms (Waterhouse et al. 2018). Because of that, we applied the BUSCO tool to quantify the completeness of each sub-genome data set in terms of the expected PCG content. The BUSCO result for the PCG in the *C. canephora* sub-genome reported 73.9 % of Complete (C) signatures (being 68.6 % Single (S) copies and 5.3 % Duplicated (D)), 5% Fragmented (F) and 21.1% Missing (M). In addition, the BUSCO result for the PCG in the *C. eugenioides* sub-genome reported 77.6% C signatures (being 72.6% S and 5% D), 4.3% F and 18.1% M. The high proportion of missing universal orthologs within sub-genomes made us wonder if their PCG compositions are different. We found that missing terms of one sub-genome are present in the other and absent for the full length PCG in the unplaced contigs available in the Cara_1.0 annotation (98% M).

The union of complete BUSCO signatures of predicted PCG in both sub-genomes sum up to 2,132 (approximately 92% of the signatures in the eudicots_odb10 reference database-version from 09/10/2020). In addition, the union of fragmented signatures from the sub-genomes summed up to 184. After filtering out overlaps between Fragmented and Complete signatures across the sub-genomes, the proportion of non-missing terms increased to 2,211 (95% of eudicots_odb10 database). Those fragmented signatures are from sequences matched with lengths below two standard deviations from the BUSCO group mean sequence length(Simão et al. 2015). In our annotation the fragmented signatures may have arisen because of our choice of only reporting a single isoform per gene. We configured AUGUSTUS to select the most likely isoform when multiple was presented. However, it is not guaranteed that the reported transcript of a given gene is the longest or even the main isoform. In addition, our choice of reporting only full-length PCG may have influenced the total number of genes evaluated during the BUSCO analyses.

Surprisingly, 327 (15.3%) complete BUSCO signatures were exclusively derived from the *C. canephora* sub-genome. Meanwhile, 413 (19.4%) BUSCO signatures were exclusively derived from the *C. eugenioides* sub-genome. When we accessed the PCG with those complete BUSCO matches that are exclusive from the *C. canephora sub-genome* (365 genes; Supplemental Table S2) and the ones that are exclusive from the *C. eugenioides* sub-genome (454 genes; Supplemental Table S2). We verified that the total number of signatures is lower compared to the total number of PCG with signatures. This difference in the number of signatures and the universal orthologous proteins is explained by the finding that some PCG was assigned to more than one BUSCO signatures and also because of the potential presence of multiple copies of some universal orthologs in a sub-genome.

Those sub-genome-specific BUSCO signatures-that are not shared with the other set of homoeologs chromosomes-can occur in a single copy (S) or multiple copies (duplicated-D) configuration. However, in some lineages, complete BUSCO signatures are more likely to be found in singletons because they are evolving under single copy control (Waterhouse et al. 2011). We propose that there is an ongoing selective pressure in *C. arabica* that constrains the number of copies of universal orthologues. Similar stronger selection constraints on the evolution of essential genes were verified in other lineages such as fungi, arthropods and vertebrates (Waterhouse et al. 2011).

### A high proportion of universal orthologs are sub-genome-specific singletons

The recent-polyploid nature of *C. arabica* gives an interesting perspective of a recently formed genome because there is a tendency of duplicated genes being silenced or removed shortly after tetraploidy formation (Buggs et al. 2012). We found that about 60% of the identified universal ortholog signatures were found to be shared by both sub genomes-they were homoeologous pairs. Homoeologous are genes in the same species that originated by speciation and were later brought back together in the same genome by allopolyploidization (Glover et al. 2016). We initially hypothesized that each sub-genome would have roughly the same number of exclusive BUSCO signatures because this gene loss would be a random process. But we verified an imbalance towards the *C. eugenioides* sub-genome keeping more universal orthologs than its *C. canephora* counterpart.

Fast and system-wide gene loss is a strategy to escape selective pressure in recently formed allopolyploids (Sankoff et al. 2010; Gaeta et al. 2007). After polyploidization, they must survive extensive genomic reprogramming to get rid of transcriptional imbalances that cause reduced fitness (Papp et al. 2003). It is possible that the verified lack of about 40% of universal orthologs-that are potentially missing homoeologs-was caused by the purifying process that benefits the retention of a single copy of some types of genes (Freeling and Thomas 2006).

The tendency for the loss of duplicated genes also drives genomes towards a simpler and more stable configuration, both structurally and in code base content (Wolfe 2001; Freeling and Thomas 2006). We presume that if the sub-genomes are segregated from each other, they would not be able to survive due the lack of fundamental molecular codebase that are better kept in single copy. It is possible that the evolutionary tendency for the following *Coffea arabica* generations is to revert back to a diploid state.

### Few GO terms of universal orthologs are exclusively enriched in the *C. canephora* sub-genome

To further elucidate the functions of those sub-genome specific universal orthologues, we analyzed their enriched GO terms. There are fourteen enriched GO terms for the *C. canephora* sub-genome exclusive BUSCO signatures (Supplemental Table 3). They are involved with biological processes (BP) such as cellular response to stress (GO:0033554, pval = 5.7E^-7^), DNA repair (GO:0006281, pval = 2.4E^-5^) and DNA recombination (GO:0006310, pval = 1.60E^-6^). Their molecular function is primarily helicase activity (GO:0004386, pval = 2.7E^-4^) and the cellular component term chromosome (GO:0005694, pval = 1.6E^-5^). We compared these fourteen enriched GO terms and we found out that ten are shared between both sub-genomes specific BUSCO signatures.

Among the enriched GO terms of sub-genome specific BUSCO signatures, there are few exclusively found in the *C. canephora* sub-genome. They are DNA repair (GO:0006281, pval = 2.4E^-5^), double-strand break repair (GO:0006302, pval = 2.5E^-4^), serine-type endopeptidase activity (GO:0004252, pval = 4.4E^-4^) and helicase activity (GO:0004386, pval = 2.7E^-4^). It is important to note that despite being exclusively enriched for BUSCO signatures within a sub-genome these BP terms are not necessarily lacking in one of the homoeolog chromosomes because thousands of genes are not universal orthologs. So, although the DNA repair and response to stress capabilities are shared between sub-genomes, we suppose that some important component of the helicase activity is coordinated by the *C. canephora* sub-genome. This sub-genome enrichment of a specific part of the universal orthologs may be a way of keeping some relevant influence over the transcriptional and genome duplication machinery. It is also possible to be a form of protection from double-strand breaks promoted by the *C. eugenioides* counterpart.

### The *C. eugenioides* sub-genome seems to be the main coordinator of gene expression in *C. arabica*

Following the trend of the total number of PCG, that is higher in the *C. eugenioides* sub-genome, the number of enriched GO terms from its exclusive BUSCO signatures is also higher compared to the *C. canephora* homoeolog. There are 167 GO terms enriched for the BUSCO signatures found exclusively in the *C. eugenioides* sub-genome (Supplemental Table S3). Ten of these 167 enriched terms are shared with *C. canephora* suggesting that *C. eugenioides* is the sub-genome with more ability of transcribing universal orthologs.

The most enriched PB term is RNA processing (GO:0006396, pval = 6.3E^-13^) followed by nuclear transport (GO:0051169, pval = 1.9E^-11^). This transport is mainly characterized as RNA export from nucleus (GO:0006405, pval = 6.0E^-7^). In addition, we also found other highly significant terms such as ncRNA processing (GO:0034470, pval = 5.9E^-6^) with its child terms tRNA processing (GO:0008033, pval = 4.9E^-5^) and rRNA processing (GO:0006364, pval = 2.4E^-3^). In addition, among the BP we could distinguish the terms mRNA processing (GO:0006397, pval = 1.0E^-5^), organelle organization (GO:0006996, pval = 1.0E^-9^) with its child terms chromosome organization (GO:0051276, pval = 2.1E^-5^) and chloroplast organization (GO:0009658, pval = 1.8E^-3^). Finally, we found the molecular function terms of RNA methyltransferase activity (GO:0008173, pval = 2.7E^-9^) and protein binding (GO:0005515, pval = 5.9E^-9^) being highly enriched as well as the cellular component terms of membrane-bounded organelle (GO:0043227, pval = 1.5E^-25^) with its child terms nucleus (GO:0005634, pval = 4.8E^-10^) and chloroplast (GO:0009507, pval = 5.7E^-23^). Taken together, these results show that an important proportion of *C. arabica*’s gene expression machinery (GO:0010467, pval = 1.70E^-5^) is controlled by the *C. eugenioides* sub-genome.

### PCG in the *C. canephora* sub-genome tend to be longer and enzymes are preferentially found in the *C. eugenioides* sub-genome

Besides the finding that there are fewer PCG encoded in the *C. canephora* sub-genome we also found that they tend to be longer than PCG encoded in the *C. eugenioides* counterpart (Supplemental Figure S2). In addition, we found that the contrasting number of PCG between the sub-genomes is even more pronounced in enzymes characterized by the InterPro scan analyses. The number of identified enzymes is always higher for the *C. eugenioides* sub-genome for every evaluated enzymatic class (Supplemental Figure S3). For example, there are 116 oxidoreductases encoded in the *C. canephora* sub-genome while 1,449 in the *C eugenioides* sub-genome. The abundance of this class of enzymes in the *C. eugenioides* sub-genome is 12.5 times higher than *C. canephora* sub-genome. A similar pattern is verified for transferases (7 times), hidrolases (8.9 times), lyases (8 times), isomerases (7.57 times), ligases (3 times) and translocases (8.84 times).

It is possible that by controlling the RNA processing and transporting machinery and, in practical terms, the gene expression (GO:0010467, pval = 1.70E^-5^), the *C. eugenioides* sub-genome are effectively causing the erosion of the gene content in *C. canephora* homoeolog chromosomes. It is also possible that a more direct-and faster-interference may be archived by TE insertions leading to pseudogenization (Yang et al. 2011). This way, the *C. eugenioides* sub-genome-with its advantage control of transcription and chromosome organization-may recruit Transposable Elements (TE) to be inserted into *C. canephora* loci. Over the past few millennia this process would render affected genes unrecognizable and explaining why there are fewer enzymes in the *C. canephora* sub-genome. In addition, this can also explain why many of the remaining genes in the *C. canephora* sub-genome are longer than their *C. eugenioides* counterpart.

### Most of the GO terms in *C. canephora* sub-genome are shared with its *C. eugenioides* counterpart, but the reciproque is not true

The difference in the number of PCG, their length, BUSCO signatures and enzymatic composition between the *C. arabica* sub-genomes made us wonder if the overall distribution of GO terms is also different between the PCG of each sub-genome, and not only a specific feature of the universal orthologs. To address this, we carefully examined the sub-genome specific BLAST2GO functional annotation results.

We found that the number of GO terms for BP is 86% higher in *C. eugenioides* sub-genome than for *C. canephora* (Figure 2A). From the 2,874 BP terms identified in *C. eugenioides* sub-genome PCG 1,414 (49%) are exclusive. Meanwhile a small proportion, only 82 (5%) of the 1,542 BP terms, are exclusively found in the *C. canephora* sub-genome (Figure 2A). Interestingly, among the exclusive BP terms from the *C. eugenioides* sub-genome we could distinguish terms such as “chromatin assembly”, “chromatin maintenance”, “chromatin organization involved in negative regulation of transcription” and “production of siRNA involved in chromatin silencing by small RNA”. Taken together, these *C. eugenioides* sub-genome specific BP terms reinforce the thesis that this sub-genome is controlling the gene expression while also controlling the chromatin organization.

**Figure 2.**
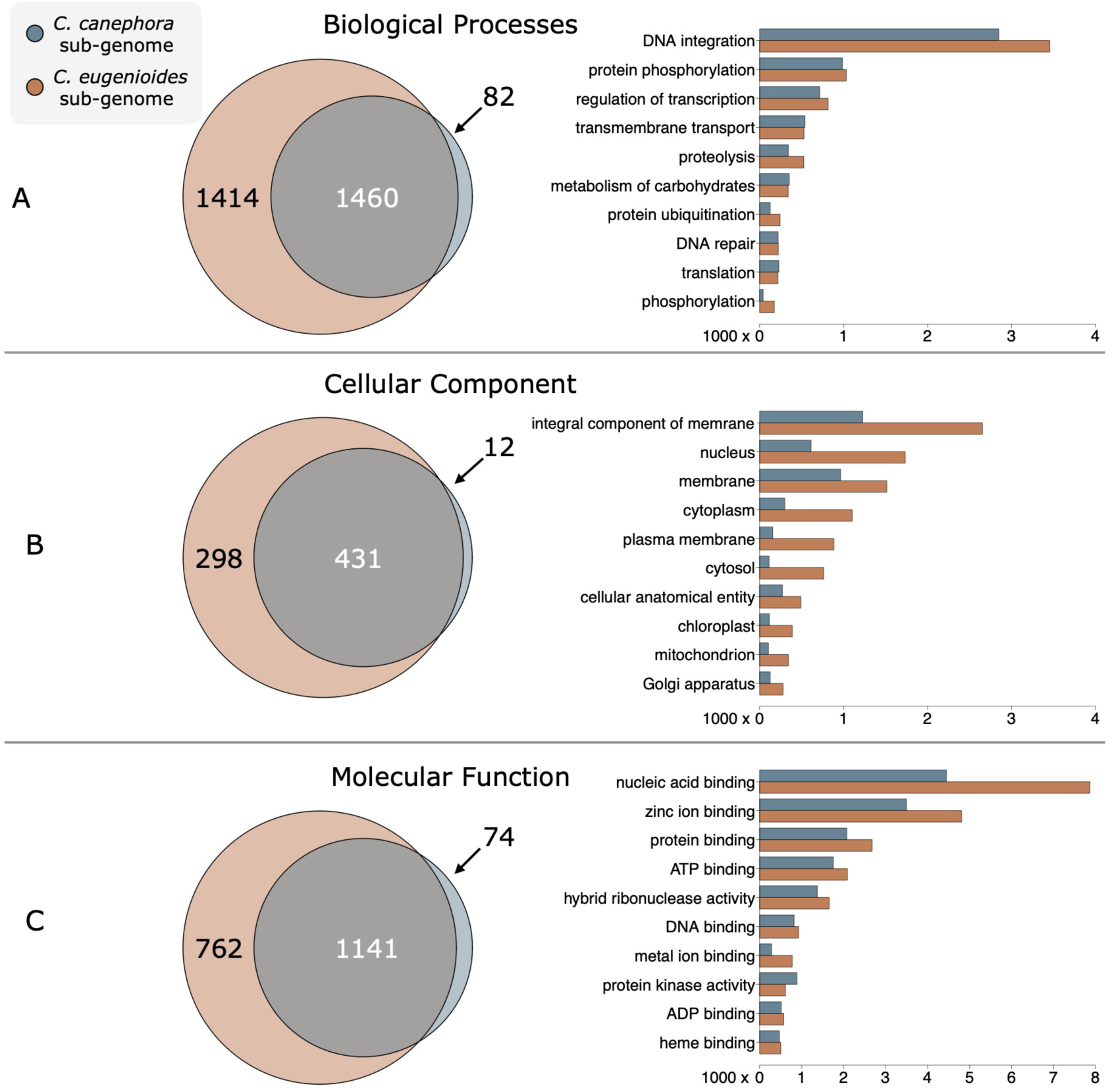
Venn diagrams of identified GO terms for sub-genome specific genes for the three main categories of Biological Processes (A), Cellular Component (B) and Molecular Function (C). Most of the GO terms were identified in the *C. eugenioides* sub-genome while a high proportion of them are not shared with its homoeologous counterpart. On the right, barplots showing a comparison of the content of PCG in each of the most abundant terms by sub-genomes. The most abundant Biological Process is DNA integration while the Molecular Function is particularly enriched for nucleic acid binding. Following the trend of total number of PCG, the *C. eugenioides* is the main contributor for all the Cellular Components, in particular the nucleus and membranes.

Regarding Cellular Component (CC) terms we found a similar pattern in which the number of individual terms is 65% higher in the *C. eugenioides* sub-genome (Figure 2B). From the 729 individual CC terms from the *C. eugenioides* sub-genome, 298 (41%) are exclusive while only twelve of 443 (3%) terms are exclusively derived from the *C. canephora* sub-genome (Figure 2B). Finally, regarding the Molecular Function (MF) terms the number of individual terms is 57% higher in the *C. eugenioides* sub-genome (Figure 2C). From the 1,903 individual MF terms from the *C. eugenioides* sub-genome, 762 (40%) are exclusive while seventy four of 1,215 (6.1%) are exclusively occurring in the *C. canephora* sub-genome (Figure 2B). We suggest that the *C. arabica* genome is still evolving towards balanced gene composition, but the *C. canephora* sub-genome suffers most of this gene loss.

### DNA integration and nucleic acid binding GO terms are dominated by Transposable Elements

In addition to the composition of GO terms being different between the sub-genomes, the number of PCG in each of the GO categories is also different. Not surprisingly, the *C. eugenioides* sub-genome is often encoding more genes within each GO category, in particular the top 10 larger categories in number of genes (bar plots at the right of 2). This outlying proportion of genes with the terms DNA integration, nucleus and nucleic acid binding led us to investigate the PCG composition of the top terms in each sub-genome (Figure 3).

**Figure 3.**
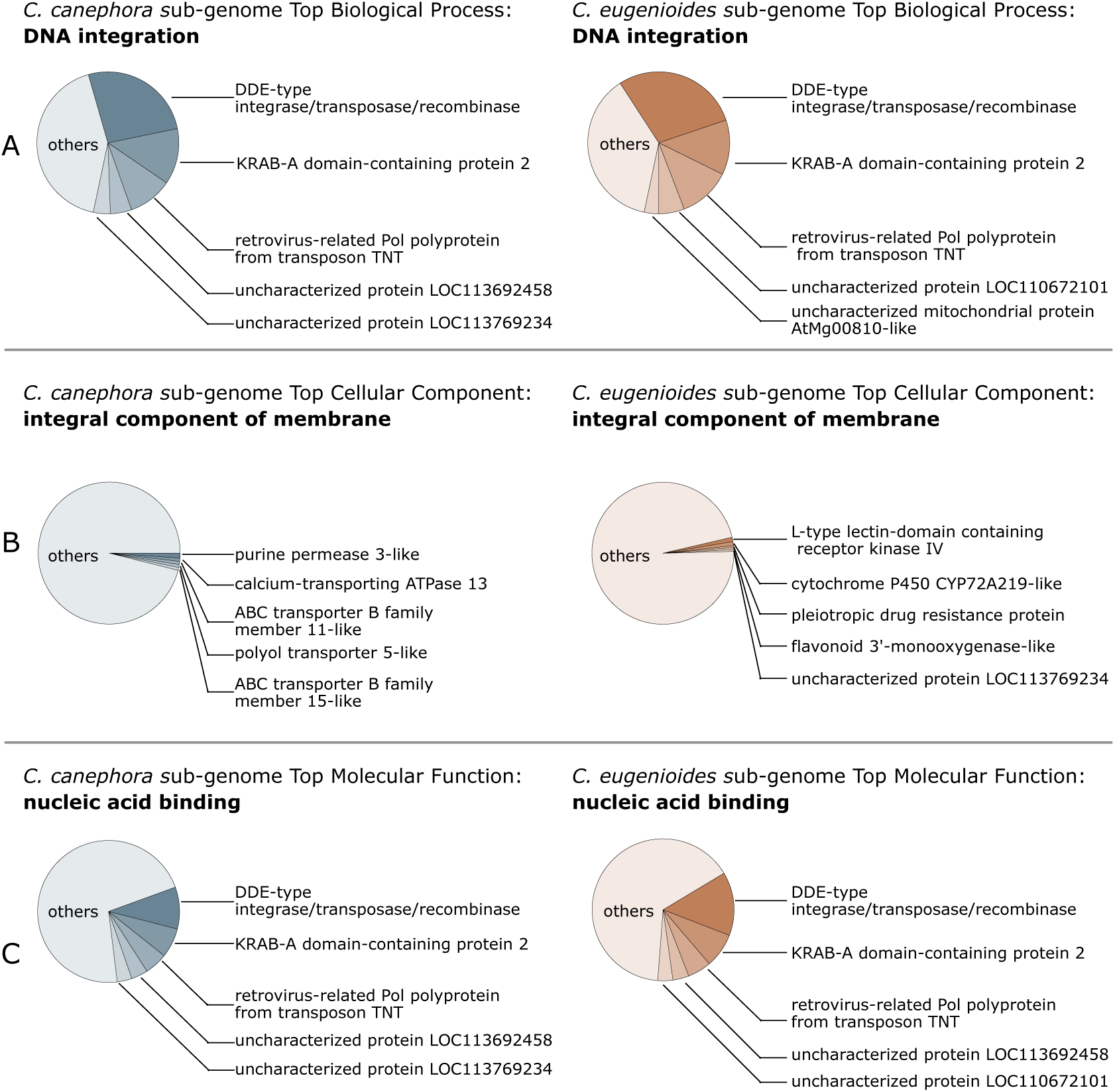
Proportion of the most abundant terms in each GO category of Biological Processes (A), Cellular Component (B) and Molecular Function (C). Known and apparent novel Transposable Elements are specially enriched in the top BP and top MF categories while the top CC category has a diverse composition. The BP term with higher number of members in both sub-genomes is “DNA integration” with 3,454 genes in the C. eugenioides sub-genome and 2,847 in the C. canephora sub-genome. The top CC “integral component of membrane” has 2,653 genes in the C. eugenioides sub-genome and 1,227 genes in the C. canephora sub-genome. The top MF “nucleic acid binding” has 7,867 genes in the C. eugenioides sub-genome and 4,447 in the C. canephora sub-genome.

This combination of abundant terms related to DNA binding made us wonder if those transcripts would be TE. In the BP category, the term “DNA integration” is mainly composed by genes identified as *DDE-TYPE INTEGRASE/TRANSPOSASE/RECOMBINASE* that accounts for approximately 28% of the PCG in this category (Figure 3A) followed by *KRAB-A DOMAIN-CONTAINING PROTEIN 2* (∼13%) and *RETROVIRUS-RELATED POL POLYPROTEIN FROM TRANSPOSON TNT* (∼11%). The other 52% of PCG in the “DNA integration” category is mostly identified as uncharacterized proteins. Many of these genes are also reported under the most enriched MF of “nucleic acid binding” where the *DDE-type* genes account for about 13%, *KRAB-A* 7% and the transposon *TNT* 5% (Figure 3C). Once Kruppel associated box (KRAB-ZFPs) are domains only present in tetrapod vertebrates (Urrutia 2003) and homology based searches to the nr database in NCBI returned TE related matches, these reported KARB-A domain containing proteins can be miss annotated TE.

Interestingly, the BP term “DNA integration” is more than 50% composed by PCG that corresponds for only five genic families and the MF term “nucleic acid binding” is following a similar pattern, with the five more abundant families accounting for about 33% of its members. On the other hand, the CC most enriched term “integral component of membrane” has a much more diverse composition (Figure 3C). The top five gene families account for less than 5% of all genes belonging to the “integral component of membrane” category while the other 789 families or individual genes are summing up the remaining 95% of membrane components. Nevertheless, the high proportion of TE in the genome composition can be verified in the second larger CC term “nucleus”. So, there is a wide-spread dissemination of TEs that seems to be a strong force shaping the evolution of *C. arabica*.

### Part of the core diurnal leaf operating system was captured while transcribing execution orders

After exploring the *C. arabica* sub-genome organization, we performed a regulatory network analysis to understand its transcriptional activation and regulatory modules related to leaf metabolism. To do so, we used 24 RNAseq libraries previously published from fully expanded leaves collected during the morning and under a heterogeneous set of environmental conditions (Cardon et al. 2022).

Our RNAseq based regulatory network inference was performed using 88,620,764 of uniquely mapped paired-end reads that aligned to approximately 51,000 loci that potentially encode PCG. These 88 million RNAseq reads were set apart from a group of 27 million of multi-mappers that could not provide locus-specific mapping resolution and potentially bias the analysis by not meeting WGCNA microarray-based assumptions. An additional filtering procedure of low expressed loci was important to exclude 34 thousand loci that accounted for only 3.7% of the transcripts. Low expressed genes have the potential of biasing the analysis by interfering with distribution assumptions and potentially incurring false-positive results due to their relatively low contribution to the transcriptome. Finally, a total of 16,610 constitutively expressed and accurately mapped putative PCG loci were evaluated.

This counting data from RNA fragments was processed under the WGCNA guidelines (Langfelder and Horvath 2008) to infer 23 modules of co-expressed genes. Module names were assigned to random colors (Supplemental Table S4). We then searched for enriched GO terms in each module and overall results are reported in the Supplemental Table S4. After that, the modules were grouped into eight clusters-named from A to H-based on the correlation of their eigengenes (Figure 4, Supplemental Dataset S1), that are individual transcripts that summarizes the overall trend in expression of any given module (Zhang and Horvath 2005).

Statistical singular enrichment analyses of GO terms associated with each module and groups of modules allowed us to unravel important biological processes coordinated by those inferred regulatory networks. This hybrid group and module-wise approach of analyses allowed us to reveal biological processes that are underlying the diurnal metabolism of *C. arabica* leaves. For example, we found that the 266 genes in group A are primarily involved in the light reactions of photosynthesis (GO:0019684, pval_Darkslateblue_ = 7.30E^-7^) and carbohydrate biosynthetic processes (GO:0016051, pval_Plum_ = 3.4E^-4^). The proteins encoded by those genes are located in the chloroplasts (GO:0009507, pval_Plum_ = 9.5E^-4^). Interestingly, a *TREHALOSE-PHOSPHATE SYNTHASE* is a key component of this group because it is the most central node, in terms of total number of connections (degree), in the Plum module.

The trehalose-6-phosphate is an important signaling metabolite that regulates carbon assimilation and sugar status in plants by balancing the synthesis and breakdown of starch (Ponnu et al. 2011). Meanwhile, the closely related group B, with 448 PCG divided in two modules, Sienna and Tan, is primarily involved with the cellular amino acid metabolic process (GO:0006520, pval_Tan_ = 2.70E^-5^). In addition, the term chloroplast (GO:0009507, pval_Tan_ = 9.7E^-4^) and others chloroplast related terms are shared between groups A and B suggesting a co-regulation between the PCG involved in light reactions of photosynthesis and cellular amino acid metabolic process in *C. arabica* leaves.

Next, 3,583 putative PCG from group C are enriched for lipid (GO:0006629, pval_Darkred_ = 7.40E^-11^), protein (GO:0019538, pval_Darkred_ = 6.70E^-8^) and carbohydrate (GO:0044723, pval_Darkred_ = 1.40E^-5^) metabolic processes-more precisely catabolic processes (GO:0009056, pval_Darkred_ = 2.00E^-7^)-in the cytoplasm (GO:0005737, pval_Darkred_ = 7.90E^-14^). The largest member of group C, the Turquoise module with 2,021 PCG components, was mostly enriched for cell wall biogenesis (GO:0042546, pval_Turquoise_ = 8.50E^-21^). In addition, the small, but significantly enriched for photosynthesis (GO:0015979, pval_coral_ = 5.10E^-12^) and thylakoid (GO:0009579, pval_Coral_ = 2.20E^-13^), Coral module was found to be directly involved with the generation of precursor metabolites and energy (GO:0006091, pval_Coral_ = 1.80E^-5^). Cell walls are dynamic structures that are constantly being maintained for enhanced structural and defense capabilities(Vaahtera et al. 2019). This group C is showing that part of the photosynthetic machinery, specifically the photosystem I (GO:0009522, pval_Coral_ = 6.80E^-11^) are in transcriptional coordination with cell wall biogenesis and catabolic processes.

Group D is the most distant in terms of their eigengenes correlation with other modules. It is composed of three modules named Orange, Greenyellow and Honeydew summing up to 695 putative PCG. Its largest module, Greenyellow, is specially enriched for protein folding (GO:0006457, pval_Greenyellow_ = 7.30E^-55^), response to heat (GO:0009408, pval_Greenyellow_ = 1.00E^-19^) and response to reactive oxygen species (GO:0000302, pval_Greenyellow_ = 8.20E^-15^). The most central gene in terms of eigengene centrality is a (chloroplastic) *SMALL HEAT SHOCK PROTEIN*. These findings allowed us to propose that this group provides response mechanisms to perceived thermic variations in the environment. This mechanism seems to be coordinated across membranes such as the ones that envelops the endoplasmic reticulum (GO:0005783, pval_Greenyellow_ = 1.40E^-7^). Changes in the membrane fluidity are used as sensors of the temperature variation (Murata and Los 1997). These signals can be transmitted to the cytoplasm from the endomembrane system (GO:0012505, pval_Greenyellow_ = 5.10E^-6^) by proteic transmembrane transporters (GO:0022857, pval_Orange_ = 4.2E^-4^). There, they can interact with protein folding mechanisms to prevent heat related damage (Mittler et al. 2012). It is possible that this group is intrinsically more reliant on the environment-more specifically the heat-and this may be the reason why these modules behave as an outgroup in the eigengene correlation cladogram (Figure 4).

**Figure 4.**
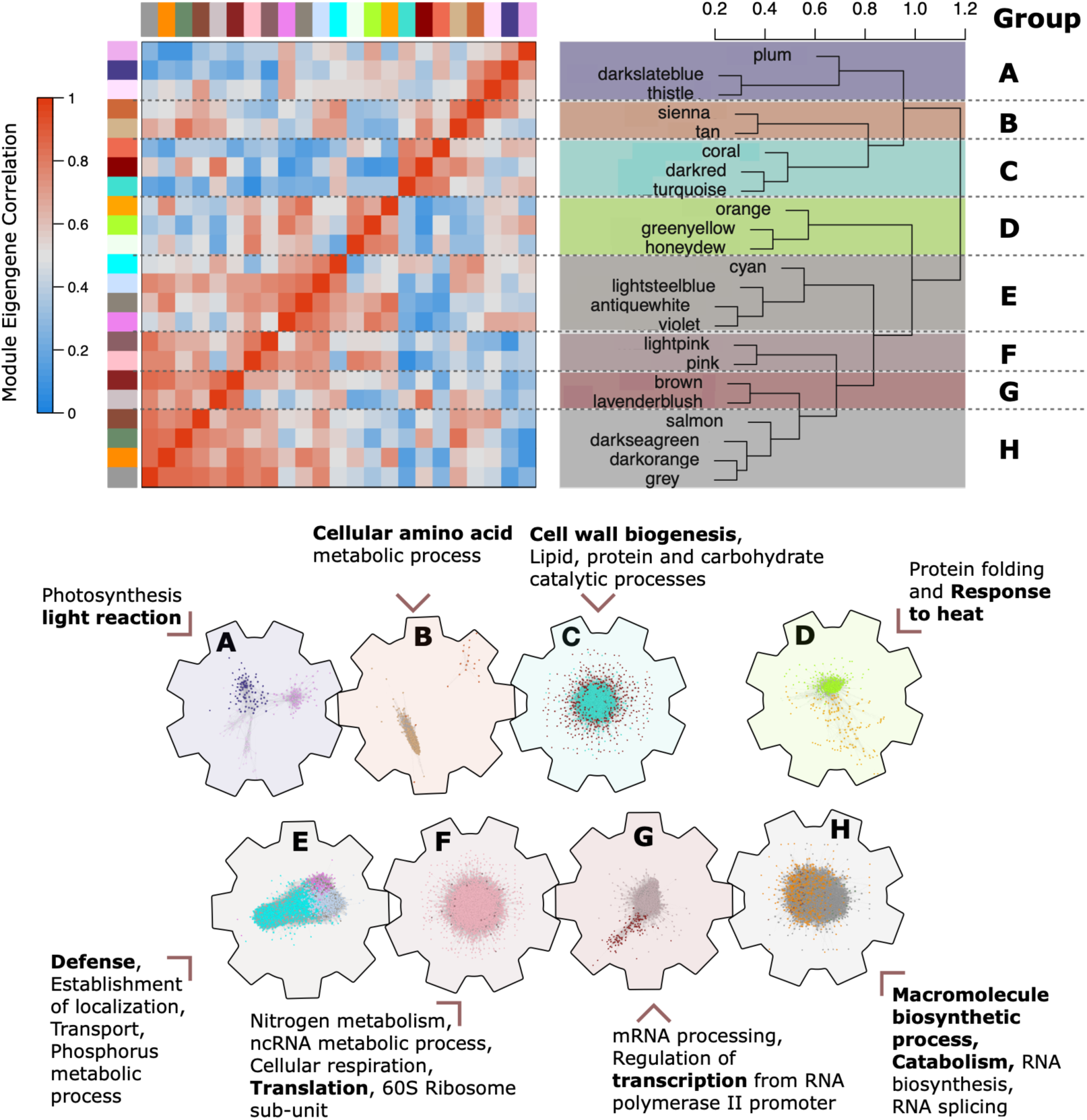
Co-expression inference revealed 23 functional modules involved with important processes of diurnal leaf metabolism. The heatmap on the left shows that the eigengenes of some functional modules are also co-expressed forming functionally larger groups. The hierarchical clustering on the top left provided delimitation for co-expression groups based on a height cutoff of 0.75. On the bottom, the schematic representation of the interconnectedness of 16,610 expressed genes. Groups of co-expressed genes, represented as engaged or disengaged gears, shows that processes such the light-reaction of photosynthesis and the amino acid and cell wall metabolism are co-regulated. Meanwhile, defense, translation, transcription and other biosynthetic processes are forming a yet larger cluster of functional modules. Some of the modules are formed by macro molecules such ribosome subunits, RNA polymerase II, spliceosomes and chromosome components.

The next group was assigned to the letter E and aggregates a total of 1,660 putative PCG that compose four modules; Cyan, Lightsteelblue, Antiquewhite and Violet. Here, we found a correlation between modules involved with establishment of localization, transport, defense and protein phosphorylation. Protein phosphorylation is an important post-translational modification that regulates protein functions and controls cellular processes during the diurnal cycle in different Arabidopsis organs and seedlings (Uhrig et al. 2019).

The enriched GO term establishment of localization (GO:0051234, pval_Cyan_ = 1.80E^-11^, pval_Lightsteelblue_ = 3.90E^-9^, pval_Antiquewhite_ = 7.2E^-4^) is shared between three module members of group E. Permeating the other enriched BP of this group is the term phosphorus metabolic process (GO:0006793, pval_Cyan_ = 2.80E^-21^, pval_Lightsteelblue_ = 6.3E^-4^). Another important phosphorus-related term is cellular response to phosphate starvation (GO:0016036, pval_Violet_ = 4.40E^-15^) that is particularly enriched in the Violet module. These phosphate starvation responses are a collective array of morphological and physiological adaptive changes that are critical for plant survival when phosphorus is limited (Chiou and Lin 2011). The physiological responses for phosphate starvation also play roles in the regulatory mechanism of phosphate usage (Chiou and Lin 2011). In addition, several inorganic phosphate (P_i_) transporters have been implicated to compose defense response mechanisms (eWang et al. 2014, 1).

Supporting the importance of phosphoric compounds to plant defense strategy is the finding that a T-DNA insertions in a loci encoding for a member of *PHOSPHATE TRANSPORTER (PHT)* family increases *Arabidopsis thaliana* susceptibility to virulent *Pseudomonas* strains (Wang et al. 2011). Here we found two members of the *PHT* family being highly connected members of the Cyan module and co-expressed with *SALICYLIC ACID-BINDING PROTEIN,* known to be involved with salicylic acid-dependent immune responses (Chan et al. 2021).

The two larger modules of group E, Cyan and Lightsteelblue with 890 and 579 members respectively, are especially enriched for membrane-related CC terms (GO:0016020, pval_Cyan_ = 1.20E^-13^, pval_Lightsteelblue_ = 4.20E^-7^). In addition, the Cyan module is particularly enriched for the BP response to salicylic acid (GO:0023051, pval_Cyan_ = 3.30E^-5^), exocytosis (GO:0006887, pval_Cyan_ = 7.70E^-7^) and golgi vesicle transport (GO:0048193, pval_Cyan_ = 1.20E^-6^). Meanwhile, Lightsteelblue module is enriched for terpenoid biosynthetic process (GO:0006721, pval_Lightsteelblue_ = 4.50E^-6^) and defense response (GO:0006952, pval_Lightsteelblue_ = 1.10E^-5^). These findings reinforce the importance of phosphorus for the immune system of plants. The members of this group E, in particular the *Cyan* module, can be further investigated to better elucidate how plants defend themselves from other organisms.

The next group was assigned to the letter F and is composed of two modules, one is the largest of all modules in this analysis and is called *Pink* with 4,847 putative PCG. The other component of this group has only 63 PCG is called *Lightpink*. Not surprisingly, Pink is also the module with more enriched GO terms, 899 of them. Meanwhile, all enriched 43 GO terms from *Lightpink* are shared with the larger module. The *Pink* module is particularly enriched for the term organo-nitrogen compound metabolic process (GO:1901564, pval_Pink_ = 1.5E^-138^). In addition, it is also significantly enriched for many other important BP terms such ncRNA metabolic process (GO:0034660, pval_Pink_ = 3.60E^-29^), rRNA processing (GO:0006364, pval_Pink_ = 3.30E^-22^), RNA splicing, via transesterification reactions (GO:0000375, pval_Pink_ = 1.20E^-14^), ligase activity, forming aminoacyl-tRNA and related compounds (GO:0016876, pval_Pink_ = 7.40E-07) and cellular respiration (GO:0045333, pval_Pink_ = 8.50E^-16^).

The co-regulation of group F is so organized that their proteins are transported to multiple intracellular components (GO:0005622, pval_Pink_ = 1.75E^-166^). Members of this groups are a relevant part of the protein composition of the cytosol (GO:0005829, pval_Pink_ = 1.9E^-60^), mitochondrial electron transport, NADH to ubiquinone (GO:0006120, pval_Pink_ = 5.80E^-5^), the full ubiquinol to cytochrome c (2.70E-09, pval_Pink_ = 2.70E^-9^) and mitochondrial respiratory chain complex III (GO:0005750, pval_Pink_ = 3.20E^-8^). They are also enriched for chloroplast thylakoid (GO:0009534, pval_Pink_ = 1.10E^-19^), nucleus (GO:0005634, pval_Pink_ = 1.10E^-23^) and apparently compose 44% of the ribosomal protein complex (GO:0005840, pval_Pink_ = 3.30E^-113^). Their components are so widespread within the cell that the few remaining CC terms not enriched in the pink module are the ones related to the extracellular part.

It is possible that the key for understanding this group-that spans most of the CC volume-relies on genes closely related to the Pink module eigengene, a 60S ribosomal protein L24-like. In fact, 277 putative PCG of the Pink module were identified as ribosomal proteins, mostly 60S ribosomal proteins. So, it is possible that the whole translational machinery (GO:0006412, pval_Pink_ = 1.20E^-134^) correlates directly to the rhythm of ribosome assembly (GO:0042255, pval_Pink_ = 9.80E^-14^) and biogenesis (GO:0042254, pval_Pink_ = 9.90E^-36^). That way, others important processes that are directly involved with translation, such as the transcriptional steps of the gene expression (GO:0006352, pval_Pink_ = 2.40E^-7^), RNA splicing (GO:0008380, pval_Pink_ = 8.20E^-16^) and organo-nitrogen compound metabolic process, such amino acids (GO:0008652, pval_Pink_ = 1.8E^-4^) and nucleotides (GO:0009117, pval_Pink_ 1.10E^-21^), are intrinsically interlinked.

The following cluster of modules is called group G and is composed of two modules summing up a total of 945 putative PCG. The main component of this group, the Lavenderblush module, is particularly enriched for GO terms relative to nucleus (GO:0005634, pval_Lavenderblush_ = 1.60E^-17^) and gene expression (GO:0010467, pval_Lavenderblush_ = 3.10E^-11^, pval_Brown_ = 2.30E^-5^). Although the “gene expression” term is also enriched in the previous group F here the descending terms differ. In this group, the influence on gene expression (GO:0010468, pval_lavenderblush_ = 8.10E^-9^) is mainly performed by the regulation of transcription from RNA polymerase II promoter (GO:0006357, pval_lavenderblush_ = 2.70E^-11^) and chromosome organization (GO:0051276, pval_lavenderblush_ = 3.70E^-6^). This group shares with group F important aspects of the gene expression process but they constitute different parts of the machinery. The importance of group G seems to be related to the transcription while group F is mostly related to the translation.

Supporting the finding that group G is an fundamental part of transcription are the enriched terms RNA helicase activity (GO:0003724, pval_Lavenderblush_ = 2.00E^-5^), histone modifications (GO:0016570, pval_Lavenderblush_ = 3.70E^-06^) such as histone methylation (GO:0016571, pval_Lavenderblush_ = 2.90E^-05^) and histone acetyltransferase complex (GO:0000123, pval_Lavenderblush_ = 5.50E^-5^) that are related to two constranting epigenetic processes that modulates the access of RNA polymerases to PCG loci. That way, it is not surprising that the eigengene of the Lavenderblush module is an ATP-dependent helicase, from the *C. canephora* sub-genome, that seems to be encompassing the transcriptional regulation of the whole group.

The final cluster of modules is group H. This group shows a coordination of expression of 4,103 genes divided into four modules. This group is mainly involved with macromolecule biosynthetic processes and is also enriched for hundreds of GO terms, 496 of them. Two modules account for about 97% of this group PCG content, the Darkorange with 1,040 and Grey with 2,930. Traditionally in the WGCNA methodology, the grey module corresponds to the set of genes which have not been clustered in any other module. Nevertheless, this is not the case in our analyses and this cluster is a bonafide module. Both Darkorange and Grey shares 121 enriched GO terms, including regulation of metabolic process (GO:0019222, pval_Darkorange_ = 3.40E^-8^, pval_Grey_ = 1.10E^-9^) and macromolecular complex (GO:0032991, pval_Darkorange_ = 4.99E^-6^, pval_Grey_ = 1.20E ^-7^). However, even sharing a significant proportion of enriched GO terms, they are also individually enriched for module-specific terms.

The Darkorange module is enriched for photosynthesis, light reaction (GO:0019684, pval_Darkorage_ = 1.10E^-6^), chlorophyll metabolic process (GO:0015994, pval_Darkorange_ = 2.00E^-5^), generation of precursor metabolites and energy (GO:0006091, pval_Darkorange_ = 3.10E^-6^). Other photosynthesis related terms are also significantly enriched. Meanwhile, Grey is particularly enriched for terms related with mRNA processing (GO:0006397, pval_Darkorange_ = 2.90E^-34^) such as mRNA splicing, via spliceosome (GO:0000398, pval_Grey_ = 7.40E^-26^) or covalent chromatin modification (GO:0016569, pval_Grey_ = 1.90E^-12^).

We noted that even sharing the enriched GO term “macromolecular complex” there are different complexes in each of the two larger modules of group H. The child terms of “macromolecular complex” in the Darkorage module are divided into photosystems I (GO:0009522, pval_Darkorage_ = 2.00E^-5^), photosystems II (GO:0009523, pval_Darkorage_ = 6.0E^-4^) and ubiquitin ligase complex (GO:0000151, pval_Darkorage_ = 4.80E^-6^). Meanwhile, “macromolecular complex” has diverse child terms in the Grey module, including the top enriched “spliceosomal complex” (GO:0005681, pval_grey_ = 6.00E^-16^) followed by other ribonucleoproteins such pre-ribosome (GO:0030684, pval_grey_ = 2.00E^-9^), small nucleolar ribonucleoprotein complex (GO:0005732, pval_grey_ = 5.6E^-4^) and Mre11 complex (GO:0030870, pval_grey_ = 1.80E^-6^). Another complexes that are formed by PCG in the Grey module include the transferase complex (GO:0061695, pval = 8.9E^-4^) that binds phosphorus-containing groups in the holoenzyme DNA-directed RNA polymerase II (GO:0016591, pval_Grey_ = 1.6E^-3^).

Members of the Grey module include components of chromosomes in the form of cohesin (GO:0008278, pval_Grey_ = 2.9E^-4^) and H4 histone acetyltransferase complex (GO:1902562, pval = 8.20E^-5^). So, we conclude that the Grey module is majoritarily involved in the transcription, partially because of its role in regulation of RNA biosynthetic process (GO:2001141, pval_Grey_ = 1.10E^-5^). Meanwhile, the Darkorange module is mainly involved with photosynthesis, more precisely the chlorophyll biosynthetic process machinery (GO:0015995, pval_Darkorange_ = 1.6E^-3^). Underlying the whole group H, biosynthetic and catabolic processes are enriched and engaged.

Finally, we also evaluated the proportion of PCG from each sub-genome in each module, however, no clear pattern emerged. Some modules were preferentially composed by PCG from a specific sub-genome, but this data does not clearly correlate with total number of genes in a module, nor the interconnectedness measured as the module density. Nevertheless, modules enriched for GO terms related to photosynthesis, transport and transcription are preferentially composed by PCG expressed from the *C. eugenioides* sub-genome. Similar results from EST of homoeolog data suggested that *C. arabica* may have specific physiological contributions derived from specific ancestors (Vidal et al. 2010). This finding supports our hypothesis that the *C. eugenioides* sub-genome is the main controller of gene expression whereas the *C. canephora* sub-genome may function as an important source of alleles to cope with the environment.

## Discussion

This system-wide approach to investigate the transcriptome of leaves of the allotetraploid *C. arabica*, a tropical perennial plant, revealed that processes such photosynthesis, cell wall biogenesis, translation, transcription, catabolism and biosynthesis are running in synchrony. Genome-wide PCG prediction shows that the two sub-genomes encode a differential set of universal orthologs. This feature may have been an important factor that allowed the establishment of this species after the allopolyploidization event.

Our evaluation of reads uniquely mapped to the sub-genomes showed that homoeologous chromosomes are, in the majority of the cases, equally used for transcription (Supplemental Figure S4). Nevertheless, we verified that in chromosomes 2 and 10 the mapped reads derived from *C. eugenioides* seem to be the majority when compared to the reads from its *C. canephora* counterpart. On a lesser level of intensity and significance, it appears that PCG in the Chromosome 6 from *C. canephora* ancestor is preferentially transcribed in comparison to its homoeologous (Supplemental Figure S4).

Other authors, using EST analyses, found that 48% of the C. arabica was transcribed from the C. canephora sub-genome and 52% were transcribed from the C. eugenioides sub-genome (Vidal et al. 2010). They also found that in 29% of 2,646 contigs had a higher contribution of one sub-genome in comparison to the other: 13% of the contigs had more ESTs from C. eugenioides sub-genome and 16% of contigs had more ESTs from C. canephora sub-genome (Vidal et al. 2010). Our results support the finding of differential sub-genome transcription. Here, by coupling PCG prediction with transcriptome sequencing, we show that chromosome organization and RNA processing PCG are preferentially encoded in the C. eugenioides sub-genome. Many of their homoeologs may have suffered processes of pseudogenization or gene deletion.

We hypothesize that by regulating the transcription and RNA transport, the *C. eugenioides* sub-genome has an edge in the decision-making regulatory networks. However, the differential contribution of homoeologous genes to the transcriptome does not necessarily correlate with genome-wide transcription levels (Mochida et al. 2004). That way, one specific homoeolog allele from a sub-genome may be preferentially expressed in a given condition, but it is unlikely, in the allotetraploid state of the *C. arabica* genome, that one sub-genome can monopolize the whole transcriptome.

Many universal orthologs-that represent the expected eudicot essential genes-are missing from the sub-genomes-especially the one inherited from the *C. canephora* ancestral parent (Figure 5). We assume that this lack of sub-genome-specific essential genes, reflected in the missing BUSCO signatures, is due to selective pressures that benefit gene expression under single copy control (Waterhouse et al. 2011). It is possible that after the origin of *C. arabica* processes of genome reorganization occurred during its evolution, ultimately causing the loss of homoeotic genes in the *C. canephora* sub-genome (Lashermes et al. 1999). It is common that after episodes of whole genome duplications multiple genes are lost due to processes of epigenetic silencing, pseudogenization and chromosome level deletions of segments containing genes (Sankoff et al. 2010).

**Figure 5.**
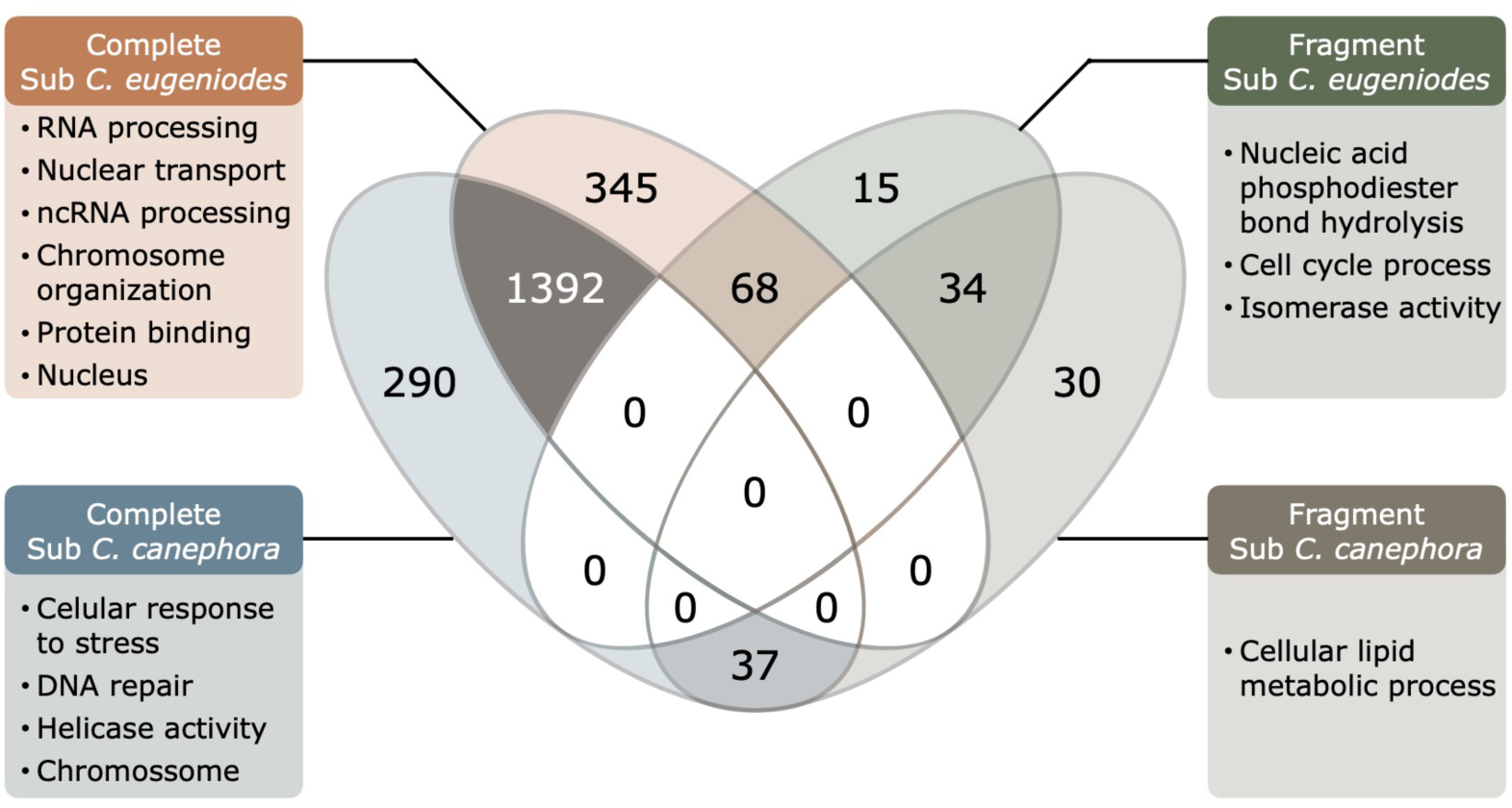
Venn diagram comparing the content of complete and fragmented BUSCO signatures of each *C. arabica* sub-genomes. About 63% (1,393) of the found BUSCO signatures are shared between both sub-genomes. A large proportion of expected universal orthologues (∼13%) were not found in the *C. canephora* sub-genome. The *C. eugenioides* sub-genome is particularly enriched for complete signatures related to RNA processing, chromosome organization and nucleus while the fewer complete signatures in the *C. canephora* sub-genome are related to responses to stresses and DNA repair.

It is yet not clear if the interspecific hybridization that created *C. arabica* had occurred in the scale of millions (Lashermes et al. 1999) or tens of thousands of years (Cenci et al. 2012; Scalabrin et al. 2020), but it is agreed that it is a recent event in evolutionary time scale (Vidal et al. 2010; Cenci et al. 2012). Recently formed polyploid species have higher extinction rates than their diploid relatives, rarely surviving over the long term and, in most cases, are evolutionary dead-ends (Arrigo and Barker 2012). Nevertheless, the few thriving individuals leave a substantial legacy in plant genomes (Arrigo and Barker 2012). Allopolyploidy can induce rapid genome evolution that may have contributed to the successful establishment of newly formed species (Ozkan et al. 2001),(Arrigo and Barker 2012).

We address that the key for allowing the survival and reproduction of *C. arabica* is the homoeologous gene loss through sequence deletion. Evidence for this hypothesis is that sequence elimination is a common and rapid phenomenon that can be verified as early as the generation F1 of newly formed hybrids (Shaked et al. 2001; Ma and Gustafson 2006). It is also possible that the missing genes in the *C. eugenioides* sub-genome turned to pseudo-genes because of our finding that gene lengths in this sub-genome are usually longer than its homoeolog counterpart. Alternatively, it is possible that the abundant PCG with GO terms related to “DNA integration”-such hundreds of *DDE-TYPE INTEGRASE/TRANSPOSASE/RECOMBINASE* in both sub genomes-are promoting ongoing insertions to *C. eugenioides* sub-genome that may be causing this gene enlargement and eventual pseudogenization.

A previous report on a set of 9,047 genes in a *C. arabica* scaffold level genome found homoeologous copies being retained in 98% of the analyzed dataset and suggested that homoeolog losses are not a random event (Lashermes et al. 2016). These authors stated that the majority of the 2% of homoeologous losses were probably due to sequence homogenization instead of sequence deletion because they verified a lack of homoeologous single nuclear polymorphisms (Lashermes et al. 2016). Our results are apparently contradicting these findings, mostly because of methodological differences. We focused our analysis on the expected homoeologous with BUSCO signatures in the *C. arabica* genome and not on a subset of genes selected by DNAseq depth-mapped to the reference genome of the relative species *C. canephora* (Denoeud et al. 2014)-and polymorphisms (Lashermes et al. 2016).

Genetic and epigenetic changes are common consequences of polyploidization in many major agricultural crop plants (Madlung and Wendel 2013), including wheat (Ozkan et al. 2001; Shaked et al. 2001), cotton(Wendel et al. 1995), rapeseed (Gaeta et al. 2007) and sugarcane (Zhang et al. 2013). Among the genetic changes verified in recent polyploids is gene loss (Buggs et al. 2012). A study with resynthesized *Brassica napus* lines-independently derived by hybridizing double haploids of *Brassica oleracea* and *Brassica rapa*-revealed the occurrence of a ‘genomic shock’ leading to gene loss in early generations following the polyploid formation (Gaeta et al. 2007). The maintained copies tend to concentrate in one of the homoeologous chromosomes (Sankoff et al. 2010). A similar rapid gene loss event may have happened early in *C. arabica* evolution and may be the reason why essential universal orthologs are missing from the sub-genomes (Figure 5).

### The gene loss in *Coffea arabica* is pronounced for homoeologous essential genes controlling gene expression

An important process in shaping genomes is purifying selection that tends to remove duplicated genes after whole genome duplications(Kondrashov et al. 2002). However, this same purifying selection causes a tendency of preservation of balanced gene activity (Bowers et al. 2003). The evolution towards balance is favored by natural selection as evidenced by the finding that angiosperms are paleopolyploid and many of them are now in a diploid state as the tetraploid genome tends to merge (Bowers et al. 2003).

The latest whole genome duplication event (the α duplication) in *Arabidopsis thaliana* happened between 83 to 86 million years ago (Bowers et al. 2003). During this geologic time, only 30% of *Arabidopsis genes* have retained syntenic copies, suggesting a large impact of gene loss on angiosperm evolution (Bowers et al. 2003). Some duplicated genes have a tendency of escaping this selective pressure whereas others are easy prey as predicted by the Gene Balance Hypothesis (Freeling and Thomas 2006). This resistance to genic duplication loss is a common evolutionary phenomenon if the duplication affects dosage dependent genes (Freeling and Thomas 2006). That way, genes which the number of its protein products are stoichiometrically calibrated to other related proteins-as the components of ribosomes-have an evolutionary tendency of keeping its duplicated copies once imbalance can cause reduced fitness (Freeling and Thomas 2006).

This balancing in the number of copies is often reflected in co-expression modules (Sankoff et al. 2010; Coate et al. 2011). The group of genes that are connected due to serialized molecular function-such enzymes in a pathway or transcriptional cascades-will be more resilient to gene loss while other groups will be more advantageous existing in single copies in the haplotype. The identified co-expression modules of *C. arabica* leaf transcriptome tended to have transcriptome components preferentially derived from one sub-genome, instead of having roughly the same number of components from each sub genome (Supplemental Table S4). It is possible that this differential module composition is a consequence of a balancing process that calibrates homoeologous copy numbers.

The machinery responsible for the regulation of transcription in *C. arabica* seems to be preferentially encoded by the *C. eugenioides* sub-genome. Module Brown, which is significantly enriched for the GO term “gene expression”, has a strong tendency of being composed by genes in the *C. eugenioides* sub-genome (65%). We initially assumed that the RNA processing machinery-the hardware component of gene expression-would mostly work as the product of dosage dependent genes. Nevertheless, the lack of essential loci related to gene expression in *C. canephora* sub-genome shows that pressures of single copy control were an important force shaping coffee evolution.

The concentration of the transcriptional decision making machinery in the *C. eugenioides* codebase may have also provided advantageous phenotypic homeostasis in response to the environment (Bertrand et al. 2015; Marques et al. 2021).The maintenance of this genomic rearrangements may have been facilitated by the preferential autogamy of *Coffea arabica*. Once it lost its self-incompatibility machinery (Lashermes et al. 1999), it was also possible that other pieces of its chromosomes were rapidly lost due to inbreeding depression.

*C. arabica* is a young species (Scalabrin et al. 2020) that seems to be evolving towards a simpler genome configuration. The stabilization of its genome left marked differences in homoeolog gene composition which was made evident by the loss of universal orthologues. In the million years to come, we expect a natural tendency of gene loss and chromosome merging. Finally, this system-wide approach clarified how biological processes (i.e., photosynthesis, cell wall biogenesis, translation, transcription, catabolism and biosynthesis) are running in synchrony. Thus, this work contributes to comprehending genome evolution of recent polyploids and supports crop breeding programs through future functional studies considering the eigengenes, unknown and/or species-specific genes found in coffee.

## Methods

### RNAseq library acquisition

Illumina® RNAseq (next-generation sequencing of complementary DNA-cDNA) samples from *C. arabica* were retrieved from the Sequence Read Archive (SRA) using the fastq-dump tool from the SRA toolkit (v. 2.9.6-1) or retrieved directly from the sequencing company when sequenced by our group. Supplemental Table S1 shows details of each Bioproject, from which approximately 174 billion nucleic bases in 81 RNAseq libraries are publicly available through SRA. Multiple coffee tissues such as beans (Cheng et al. 2020; Cheng et al. 2018), leaves (de Oliveira et al. 2020; Cardon et al. 2022), seeds (Stavrinides et al. 2020) and roots (dos Santos et al. 2019) were used to find exons and predict genes in this study.

### Library Quality Control

RNAseq libraries were inspected for adapters using the minion tool from the kraken package (Davis et al. 2013). Afterwards, adapters and low quality reads were processed with Trimmomatic v. 0.39 (Bolger et al. 2014) using the parameters ILLUMINACLIP:3:25:6, SLIDINGWINDOW:4:20 and MINLEN:30. All the quality-controlled reads in each group from Supplemental Table S1 were used for a *de novo* transcriptome assembly and genome mapping.

### Generating training Gene Structures from Short Read RNAseq Data

The steps described in this section are in accordance with the *Basic Protocol 1* and *Support protocol 2* from “Predicting genes in single genomes with AUGUSTUS” (Hoff and Stanke 2019). All the quality-controlled reads were mapped to the *C. arabica* Caturra genome available at NCBI (BioProject PRJNA506972) (Johns Hopkins University 2018). The fasta sequence of the complete genome was retrieved and used as reference to map the reads of all libraries in Supplemental Table S1 using hisat2 v. 2.1.0 with default parameters (Kim et al. 2019, 2). After that, the samtools package v. 1.10 (Li et al. 2009) was used to convert sam to bam files, remove all unmapped reads, remove all multi-mappers and filter out paired-end reads with unmatching pairs. Then, picard tools (2019) was used to remove read duplication. Finally, all the libraries were combined in a single bam file with the samtools merge command.

Uniquely mapped paired-end *RNAseq* fragments were then subdivided according to which sub-genome they mapped using samtools and were sorted based on their chromosome coordinates, spamming a length of 989.98 million bases. All the subsequent steps of AUGUSTUS training and prediction were run separately for each of the *C. arabica* sub-genomes. The unplaced contigs, with 104.37 million nucleotides, were not evaluated mostly because it consists of highly repetitive sequences with low PCG density. Based on the annotation Cara_1.0 it is estimated that the unplaced contigs encode approximately 13 PCG per million bases while the assembled chromosomes presented an average of 65 PCG per million bases (Supplemental Table S5).

To reduce potential noise due to coincidental alignments a filtering step was applied with “filterBam” from the AUGUSTUS package with parameters “unique”, “paired” and “pairwiseAlignment”. Next, an additional sorting step was applied. Intron information was retrieved with the program “bam2hints” with the parameter “intronsonly”. Once most of the retrieved RNAseq data was unstranded – it is not known from which strand a fragment originated – the script “filterIntronsFindStrand.pl” was applied to identify the correct strand by using genomic splice site information. In this step all reads that did not have appropriate splice site information were removed.

Template genes for training AUGUSTUS were generated with unsupervised GeneMark-ET (Hoff et al. 2016) training procedures using the script “gmes_petap.pl” with the intron information retrieved in the previous step. This *ab initio* prediction was then filtered with the script “filterGenemark.pl” to select putative genes that have support in the RNA-seq alignment. Because AUGUSTUS requires information of both the coding and non-coding sequences, the flanking region of the *ab initio* predicted genes was evaluated. The script “computeFlankingRegion.pl’’ was used to calculate the average length of genes and the flanking region was set to 230 for both sub-genomes (approximately half of the average mRNA size) in the script “gff2gbSmallDNA.pl”. The resulting file was processed in accordance with *protocol 2* in Hoff and Stanke (2019) to remove redundant gene structures at the amino acid level. This step was performed to avoid overfitting the gene prediction model. For that reason, no two *ab initio* predictions of PCG with more than 80% of similarity in the primary structure were allowed in the training dataset. The resulting file was used during the AUGUSTUS prediction as hints for the identification of exons, introns and UTRs.

### *De novo* Transcriptome Assembly and protein coding transcripts inference

All the quality-controlled reads retrieved from libraries that shared the same description (Supplemental Table S1) were submitted to Trinity v. 2.8.5 (Grabherr et al. 2011) to perform the *de novo* assembly of the transcriptome in each evaluated condition. The used parameters for each run were “seqType fq”, “CPU 24”, “full_cleanup”, “max_memory 140G”, “min_contig_length 50”, and “no_normalize_reads”. Statistics about each assembly were retrieved using the “TrinityStats.pl” script and, to remove excessive isoforms and variants for each transcripts sequences with 95% of similarity were collapsed using cd-hit-est v. 4.8.1 (Li and Godzik 2006) with parameters “c 0.95”, “n 10”, “T 0” and “M 0”.

For each of the remaining putative genes in all the assembled transcriptomes only the largest isoform was selected with the script “get_longest_isoform_seq_per_trinity_gene.pl”. This was performed to avoid the effect of multiple isoforms of long genes to influence the average contig length and also to speed up the further prediction steps. Additionally, all the predicted transcriptomes from *de novo* assemblies were combined in a single file and sequences with more than 80% of similarity were collapsed with cd-hit-est v. 4.8.1 (Li and Godzik 2006) with parameters “c 0.80”, “n 5”, “T 0” and “M 0”.

Next, the longest Open Reading Frame (ORF) for each transcript was inferred with Transdecoder.LongOrfs with parameter “m 16” which reflects nucleotide sequences of 48 bp. This was performed to allow a lower limit able to include short proteins that are more likely to be orphan genes. Then, the program TransDecoder.Predict v. 5.5.0 was run with the parameter “single_best_only” to select the most probable protein coding transcripts of a given genomic locus. Finally, the software Benchmarking Universal Single Copy Orthologs (BUSCO) v. 4.1.4 (Simão et al. 2015) with parameters “l eudicots_odb10”, “m transcriptome” and “c 24” was used to evaluate the transcriptome based protein coding inference (Simão et al. 2015).

### Generating training gene structures from proteins

The steps described in this section are in accordance with the *Alternate protocol 1* from “Predicting genes in single genomes with AUGUSTUS” (Hoff and Stanke 2019). The pipeline was run separately for each sub-genome. Firstly the predicted protein sequences were mapped to the sub-genomes using GenomeThreader v 1.7.1 (Gremme et al. 2005) through the encapsulating script “startAlign.pl” from the Braker software (Hoff et al. 2016). Then, the resulting alignments were converted to the Gene Transfer Format (GTF) with “gth2gtf.pl” script. Next, the flaking region length was computed with “computeFlankingRegion.pl” and was set to 978 and 1001 to *C. canephora* and *C. eugenioides* sub-genomes respectively. Those values were then used to generate a GenBank flat file with “gff2gbSmallDNA.pl’’ required in the subsequent training steps.

### Generating training gene structures from mRNA

The steps described in this section are in accordance with the *Alternate protocol 2* and *support protocol 5* from “Predicting genes in single genomes with AUGUSTUS” (Hoff and Stanke 2019). The pipeline was run separately for each sub-genome. The putative nucleotide sequences (CDSs) of protein coding loci identified by Transdecoder were used as a proxy for Expression Sequence Tags (EST) to help improve the prediction of exons, introns and UTR regions. To do so the Program to Assemble Spliced Alignments (PASA) v. 2.4.1 (Haas et al. 2008) was used. Firstly, the transcripts were cleaned using seqclean. Then, a configuration file was created with the parameters “MIN_PERCENT_ALIGNED=0.8”, “MIN_AVG_PER_ID=0.9” and “m=50”. The PASA pipeline was called with the parameters “C”, “R”, “CPU 8” and ‘ALIGNERS blat”. The alignment was performed with the blat tool v. 35×9 (Kent 2002). Next, the ORFs were calculated with “pasa_asmbls_to_training_set.dbi” and incomplete ORFs were filtered out with custom scripts to create *a bonafide* file. Next, the flanking region length was computed with “computeFlankingRegion.pl” and was set to 1,323 and 1,343 to *C. canephora* and *C. eugenioides* sub-genomes respectively.

### Protein coding gene prediction using extrinsic evidence

The steps described in this section are in accordance with Basic *protocol 3, alternate protocol 7* and *alternate protocol 8* from “Predicting genes in single genomes with AUGUSTUS” (Hoff and Stanke 2019). The pipeline was run separately for each sub-genome. Firstly, we generated hints using the paired-end *RNAseq* alignments produced in the section “Generating training Gene Structures From Short Read RNAseq Data**”.** That information was helpful because they encoded the probable locations of introns and the coverage of transcribed exons and UTRs loci. Complementary to the intron hints were generated in *Basic protocol 1*, the exon hints where produced with “bam2wig” and them processed with the script “wig2hints.pl” with parameters “width=10”, “margin=10”, “minthresh=2”, “minscore=4”, “prune=0.1”, “src=W”, “type=ep”, “UCSC=unstranded.track”, “radius=4.5”, “pri=4” and “strand="."”.

Next, we generated hints from the protein data that can aid the prediction of CDS, introns, the correct reading frame and the position of start and stop codons. The result of the alignment with GenomeThreader (performed previously) was used as input in the script “align2hints.pl” with default parameters. Next, *Alternate Protocol 8* (Hoff and Stanke 2019) was adapted to generate hints from *de novo* PCG predicted from Trinity and Transdecoder. Only complete sequences (with 5’ UTR, CDS and 3’ UTRs) were considered. The BLAT v. 36×9 (Kent 2002) was used to align those sequences to the sub-genomes with the parameters “noHead” and “minIdentity=92”. Then, “pslCDnaFilter” was applied to filter the potentially most useful alignments with parameters “minId=0.9”, “localNearBest=0.005”, “ignoreNs” and “bestOverlap”. The hints file was then produced with “blat2hints.pl” with parameters "minintronlen=35” and “trunkSS”. Finally, all the hints from the different sources of evidence (such as proteins, cdna and RNA-seq) were combined in a single file to guide AUGUSTUS during PCG prediction.

### Training Augustus for specie specific parameters

During all the steps of prediction of *ab initio* gene structures, the program “etraining” was run to create and/or update species-specific parameters of gene models. Each of the “etraining” runs were performed on a set of approximatively 80% of the putative gene structures and then tested in a subset containing the remaining 20%. A final training step was performed by running the script “optimize_augustus.pl”. In addition, to increase the accuracy of AUGUSTUS gene prediction and to identify potential regulatory loci in the optimization procedure of *C. arabica*, PCG were performed with the UTR training steps described in the *support protocol 5* (Hoff and Stanke 2019).

### Running AUGUSTUS with hints and PCG annotation

After successive stages of model training and testing for both sub-genomes the AUGUSTUS was finally run in accordance with the *Basic Protocol 4* from “Predicting genes in single genomes with AUGUSTUS” (Hoff and Stanke 2019). The extrinsic support from multiple sources were combined and the parameters set to “UTR=on”, “allow_hinted_splicesites=atac”, and “genemodel=complete”. Only one transcript was allowed per gene, i.e., no multiple isoforms were allowed to be reported.

The annotation of PCG was performed using blast2GO (Götz et al. 2008) by homology searches with blastp (Altschul et al. 1997) against the RefSeq protein database (O’Leary et al. 2016). In addition, functional analysis was also performed with InterProScan by classifying the predicted proteins into families and identification of domains and important functional sites (Blum et al. 2021). Then, Gene Ontology (GO) terms were mapped and processed for each putative gene with the blast2GO annotation tool. That way, an annotation rule was applied to the found ontology terms for each putative gene. This rule was set to default parameters with the aim of finding the most specific annotations within a certain level of reliability. Finally, the BUSCO software v. 4.1.4 (Simão et al. 2015) was run with the predicted protein sequences from each sub-genome and parameters set to “l eudicots_odb10”, “m prot”, “--long” and “c 24” to evaluate the completeness of the protein coding inference (Simão et al. 2015).

### Co-expression network analysis of PCG on fully expanded leaves

To infer co-expression modules (clusters) of expressed PCG in fully expanded leaves of *C. arabica* we applied procedures from the Weighted Gene Co-expression Network Analysis (WGNA) R-package (Langfelder and Horvath 2008). The libraries from the bioproject ID PRJNA851465 were selected because they represent leaves under a heterogeneous set of environmental conditions which are expected to trigger multiple regulatory networks to cope with the field variability (Cardon et al. 2022). The BioSamples covered two cities (Pirapora and Varginha) during two harvest times (April and October) and with two *C. arabica* cultivars (Acauã and Catuaí). After *inhouse* quality processing steps, the RNAseq reads were mapped to the *C. arabica* reference genome (BioProject accession PRJNA506972) (Johns Hopkins University 2018) using the STAR aligner v. 2.7.8 (Dobin et al. 2013). Then, fragments mapped to the gene exons of our prediction were quantified with the HTseq-count script (Anders et al. 2015) and processed with required transformations that met the requirements for the WGNA.

First, PCG with mean expression below 25 counts per library were filtered out. Next, we normalized the count data in Counts Per Million (CPM) and then we log2 transformed the matrix of CPM values to fit the assumptions of the WGCNA package. After that, we calculated an adjacency matrix of Pearson correlations between all pairs of expressed loci and raised it to a power **β** (soft threshold) of 6. The **β**=6 parameter was based on the scale free topology criterion (Zhang and Horvath 2005). After that, to minimize the effect of noise and spurious associations, we transformed the adjacency matrix into a Topological Overlap Matrix (TOM). Next, a dendrogram, with the co-expression modules as its branches, was inferred based on the average dissimilarity of the TOM using the Dynamic Tree Cut method. Then, we analyzed the individual co-expression modules with the R package igraph (Csardi and Nepusz 2006). Finally, the GO terms for all members of each module were analyzed with the web tool agrigo2 (Yan et al. 2017) to investigate for enriched terms that provided clues about the biological processes, localization and molecular function of these co-expressed genes. The statistical analysis was performed using the singular enrichment model and a significance cutoff was set using the Benjamini and Hochberg false discovery rate (Benjamini and Hochberg 1995) of 0.05.

## Competing Interest statement

The authors declare the following financial interests/personal relationships which may be considered as potential competing interests:

Antonio Chalfun Junior reports financial support was provided by the Brazilian National Institute of Science and Technology for Coffee, Minas Gerais State Foundation of Support to the Research. and Brazilian National Council for Scientific and Technological Development.

## Data access

All data used in this study were retrieved from SRA public databases under the BioProject accession numbers PRJEB24850, PRJEB24137, PRJNA609253, PRJEB32533, PRJEB15539, PRJNA851465. The *C. arabica* Caturra genome was retrieved from BioProject PRJNA506972. The predicted putative PCG sequences, coordinates and annotation are available at https://dbi.ufla.br/lfmp/ca_annotation. Source codes for all the analysis are available at GitHub https://github.com/thalescherubino/thesisChapter1

## Acknowledgments

We thank Blake C. Meyers for comments on the manuscript.

